# Crosstalk between Ethylene and ABA during changes in soil water content reveals a role of ACC in coffee anthesis regulation

**DOI:** 10.1101/2021.08.10.455871

**Authors:** Marlon Enrique López, Iasminy Silva Santos, Robert Márquez Gutiérrez, Andrea Jaramillo Mesa, Carlos Henrique Cardon, Juliana Maria Espindola Lima, André Almeida Lima, Antonio Chalfun-Junior

## Abstract

- Coffee (*Coffea arabica* L.) presents an asynchronous flowering regulated by endogenous and environmental stimulus, and anthesis occurs once plants are rehydrated after a period of water deficit.
- We evaluated the evolution of Abscisic Acid (ABA), ethylene, 1-aminocyclopropane-1-carboxylate (ACC) content, ACC oxidase (ACO) activity, and expression analysis of the *Lysine Histidine Transporter* 1 (*LHT1)* transporter, in roots, leaves and, flower buds from three coffee genotypes (*Coffea arabica* L. cv Oeiras, Acauã, and *Semperflorens*) cultivated under field conditions with two experiments. In a third field experiment, the effect of exogenous supply of ACC in coffee anthesis was evaluated.
- We found an increased ACC level in all tissues from the three coffee genotypes in the re-watering period just before anthesis for all tissues and high expression of the *LHT1* gene in flower buds and leaves. Ethylene content and ACO activity decreased from rainy to dry period whereas ABA content increased. Higher number of opened and G6 stage flower buds were observed in the treatment with exogenous ACC.
- The results showed that the interaction of ABA-ACO-ethylene and intercellular ACC transport among leaves, buds, and roots in coffee favors an increased level of ACC that is most likely, involved as a modulator in coffee anthesis.

## 1. INTRODUCTION

Worldwide, the production of most crops depends on flowering, and particularly in some species, such as coffee (*Coffea arabica* L.), this process can directly influence the quality of the final product (cup quality). Coffee asynchronous flower bud development leads to uneven flowering (De Oliveira *et al*., 2014) and, depending on the geographical location of the plantation, it can result in various flowering events. Flowering is an important step of the coffee reproductive phase, being influenced by different endogenous and environmental signals. In this process, one of the most important elements is water as a signaling transporter, as it has been shown that coffee anthesis occurs after rain events preceded by a period of moderate water stress (Drinnan & Menzel, 1994). Other factors, such as photoperiod, temperature, shade conditions, plant nutritional status, and phytohormones can also affect floral transition and plant development (De Camargo, 1985; Ramírez *et al*., 2010).

Information on the involvement of plant hormones in coffee flowering regulation is scarce. Previous studies have shown that Abscisic Acid (ABA) increases during the dry period and is associated with coffee flower bud dormant or “latent state”, having plant rehydration decreasing ABA and increasing Gibberellin (GA) levels (Cueto & Dathe, 1986). After water withdrawing stress, dormant and non-dormant G4 flower buds (ranging from 3.2 to 6 mm in length) (Morais *et al*., 2008) could be distinguished based on their ethylene evolution (Schuch *et al*., 1992). In other species, changes in ethylene levels can either delay or promote flowering, as observed in Rice (*Oryza sativa* L.) (Wang *et al*., 2013), Arabidopsis (*Arabidopsis thaliana*) (Achard *et al*., 2007), Pineapple (A*nanas comusus* L.) (Trusov & Botella, 2006), and Roses (Meng *et al*., 2014).

The relationship between ABA and ethylene in the regulation of different biological processes is well known. Similar to ethylene, ABA accumulation accelerates the senescence of cut flowers and flowering in potted plants (Liu *et al*., 2014). Exogenous applications of ABA in Roses (*Rosa hybrida* L.) increased the ethylene sensitivity as well as the expression of some ethylene receptors (Müller *et al*., 2000). In hibiscus (*Hibiscus rosa-sinencis* L.), ABA negatively regulates the expression of the ethylene biosynthesis genes during flower development (Trivellini *et al*., 2011). On the other hand, the exogenous application of ABA positively regulates ethylene biosynthesis, increasing the ACC content in the abscission zone of the Lupine flower (*Lupinus luteus* L.) (Wilmowicz *et al*., 2016).

Recently, it was proposed that ethylene can play an important role in coffee flowering, once rehydrating droughted plants can increase ethylene levels and ethylene sensitivity, regulating coffee anthesis (Lima *et al*., 2020). Ethylene is a volatile compound that can be easily transported by diffusion throughout cells and intercellular spaces (Alonso & Ecker, 2001). Ethylene is produced from the amino acid methionine, which is first converted into S-adenosylmethionine (S-AdoMet) through the S-AdoMet synthetase enzyme, and subsequently, 1-aminocyclopropane-1-carboxylate (ACC) synthase (ACS) converts S-AdoMet into 1-Aminocyclopropane-1-Carboxylic Acid (ACC), that is considered the limiting step of the ethylene biosynthesis pathway. Finally, ACC oxidase (ACO) generates ethylene by oxidizing ACC (Yang & Hoffman, 1984).

The role of ACC, as a signaling molecule independent of ethylene biosynthesis, has been reviewed, showing that ACC is part of many biological processes in plants such as in cell wall metabolism, vegetative development, stomatal development, and pollen tube attraction (De Poel & Van Der Straeten, 2014; Vanderstraeten & Van Der Straeten, 2017; Polko & Kieber, 2019; Mou *et al*., 2020). In addition, it was also found that ACC has a distinct function than ethylene in the non-seed plant *Marchantia polymorpha* L. (Li *et al*., 2020; Katayose *et al*., 2021), and in the induction of sexual reproduction of the Marine red alga (*Pyropia yezoensis)* (Rodophyta) (Uji *et al*., 2020).

These different biological functions attributed to ACC seem to have originated an evolutionarily conserved signal that predates its efficient conversion to ethylene in higher plants (Li *et al*., 2020). ACC transport throughout the plant occurs by two amino acid transporters, LHT1 and LHT2 (Shin *et al*., 2015; Choi *et al*., 2019), having the *LHT1* higher activity described in root tissues and flower development (Mou *et al*., 2020). In light of this knowledge, research on the role of ACC associated with the development and growth of plants should be expanded, including the reconsideration of physiological processes primarily attributed to ethylene.

The main objective of this research was to evaluate the modulation activity of ACC in the coffee flowering process quantifying the evolution of ABA, ACO, ACC, and ethylene in the most critical development period for coffee flower buds, including flower bud development, bud dormancy, and anthesis; and the *LHT1* gene expression evaluation as an ACC intracellular transporter. It is possible that ACC, in addition to be the ethylene precursor, may be involved in the regulation of anthesis and fertilization of the coffee flower. To corroborate this hypothesis, field experiments were carried out complemented with laboratory analysis.

## 2. MATERIALS AND METHODS

To elucidate the modulation of ACC in coffee anthesis, three field experiments were designed. In each experiment, we sought to answer three complementary hypotheses related to hormonal crosstalk, the role of water, and the response of coffee plants in the exogenous application of ACC in the coffee anthesis process (Supporting information Table S1).

### 2.1 Field experiment I

To evaluate the evolution of phytohormones (ABA, Ethylene), intermediaries in the ethylene biosynthesis pathway (ACC and ACO), and relative expression of *LHT1* gene in coffee flowering development, three coffee genotypes (*Coffea arabica* L.) were selected: Oeiras, Acauã, and *Semperflorens*, classified as early, late and continuous, respectively, regarding their flowering and fruit ripening pattern. The experiment was conducted in a five-year-old coffee plantation at the Department of Agriculture of the Federal University of Lavras (UFLA) following a randomized block design with three biologicals repetitions, each one comprising ten plants. The experiment was carried out from May to August 2020, with samplings happening at the end of the rainy period (May 18th, late fall), end of dry period (August 18th, winter), and re-watering period (August 28th, late winter, 10 days after the first rain event). For each period, sampling was comprising of roots (15 to 25 cm deep into the soil), leaves (young and fully expanded at the third or fourth node from plagiotropic branches), and flower buds (G2 buds with a broad and flat apex, G3 buds up to 3 mm in length, and G4 buds ranging from 3.2 to 6 mm in length) (Morais *et al*., 2008), taking place from 8:00 till 10:00 am. Samples were immediately frozen in liquid nitrogen and stored at -80 °C to be evaluated for ACC and ABA content, ACO activity, and *LHT1* expression analysis. For the ethylene analysis, leaves, buds, and root samples were collected in glass tubes. Plant water status was assessed by measuring predawn leaf water potential (from 03:30 till 05:30 am) using a Scholander-type pressure chamber (Fig. 1a). Both quantifications were made according to Lima *et al*. (2020).

**Fig. 1.**
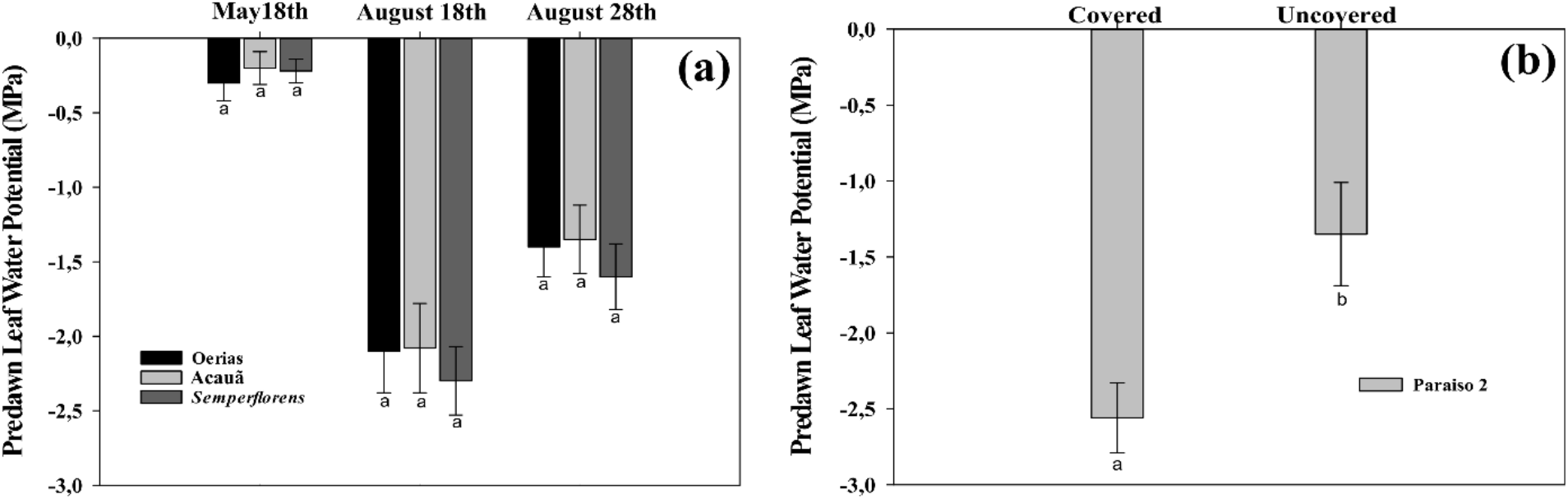
Predawn leaf water potential for coffee genotypes. **a**) Coffee plants from field experiment I in the rainy period (May 18^th^), dry period (August 20^th^), and re-watering period (August 28^th^) for Oeiras, Acauã, and *Semperflorens* genotypes **b**) Coffee plants from field experiment II in the re-watering period (August 28^th^) for Paraiso 2 coffee cultivar. Data are the mean ± 95 % of Standard Deviation (SD) of the mean (n = 8). Different letters within the same period indicate statistical (P *<* 0.05) differences within the same period, respectively.

### 2.2 Field Experiment II

Aiming to evaluate the effect of rain and hormonal balance in coffee flowering corroborating the results of the field experiment I, a second experiment was conducted in a three-year-old coffee (*Coffea arabica* cv. Paraiso 2) plantation. The experiment consisted of the evaluation of two treatments, plants under normal field conditions (Uncovered) and plants under rainfall exclusion (Covered), with three biologicals repetitions, each one composed of 10 plants in each treatment. Rainfall exclusion was achieved by the installation of translucent nylon (polypropylene) fixed on twelve 3.2 m high eucalyptus wood logs, deep 0.60 m in the soil with three wood logs 3 m distant from each other. Every eucalyptus wood log was arranged on each posterior side and in the middle of the coffee line. A translucent nylon piece of 0,2 micrometers was fixed on top of wood logs using a wooden piece of 5×7 cm with a small hobnail, covering a total of 81 m^2^ of rain exclusion area (Supporting information Fig. S1). Leaf water potential measurement (Fig. 1b) and tissue sampling for the evaluation of Ethylene, ACC and ABA content, ACO activity, and *LHT1* expression analysis were performed similarly to field experiment I.

### 2.3 Field experiment III

To better understand the effect of ACC modulating coffee anthesis, exogenous treatments of ACC were evaluated. The experiment was carried out in the coffee germplasm collection of the experimental area of the Department of Agriculture (UFLA), on adult, four-year-old coffee cultivar (*Coffea arabica cv. Semperflorens*) trees. The experiment was conducted in a randomized design with four treatments (One plant per treatment) and ten replicates per treatment (Ten branches per plant) with 5-6 nodes per branch. Four treatments were applied, ACC (300 mMol), 1-Methylcyclopropene (1-MCP, 50 mg a.i. L^-1^), ACC+1-MCP, and a control (water). It was added to all treatments a surfactant (Tween 20^®,^ at 0,08%) to improve the solutions adherence and penetration in the coffee tissues; 200 mL of total solution per treatment was used during applications. ACC concentration applied was determined according to the ACC content found in coffee tissues in the re-watering period of field experiment I (Fig. 2(1c)). 1-MCP concentration applied was determined according to Lima *et al*. (2020). Each treatment was sprayed carefully in each branch, totally wetting leaves and flower buds in the upper third of the plant. Three days after applications, a rain event of 35 mm occurred, having all treatments influenced by the same quantity of water in the field. Flower bud differentiation was evaluated fifteen days after treatment application (Previous to anthesis) counting the number of flower buds in G2, G3, G4, and G6 stages (Morais *et al*., 2008) from all nodes (5-6) in each branch (240-250 flower buds per treatment).

**Fig. 2.**
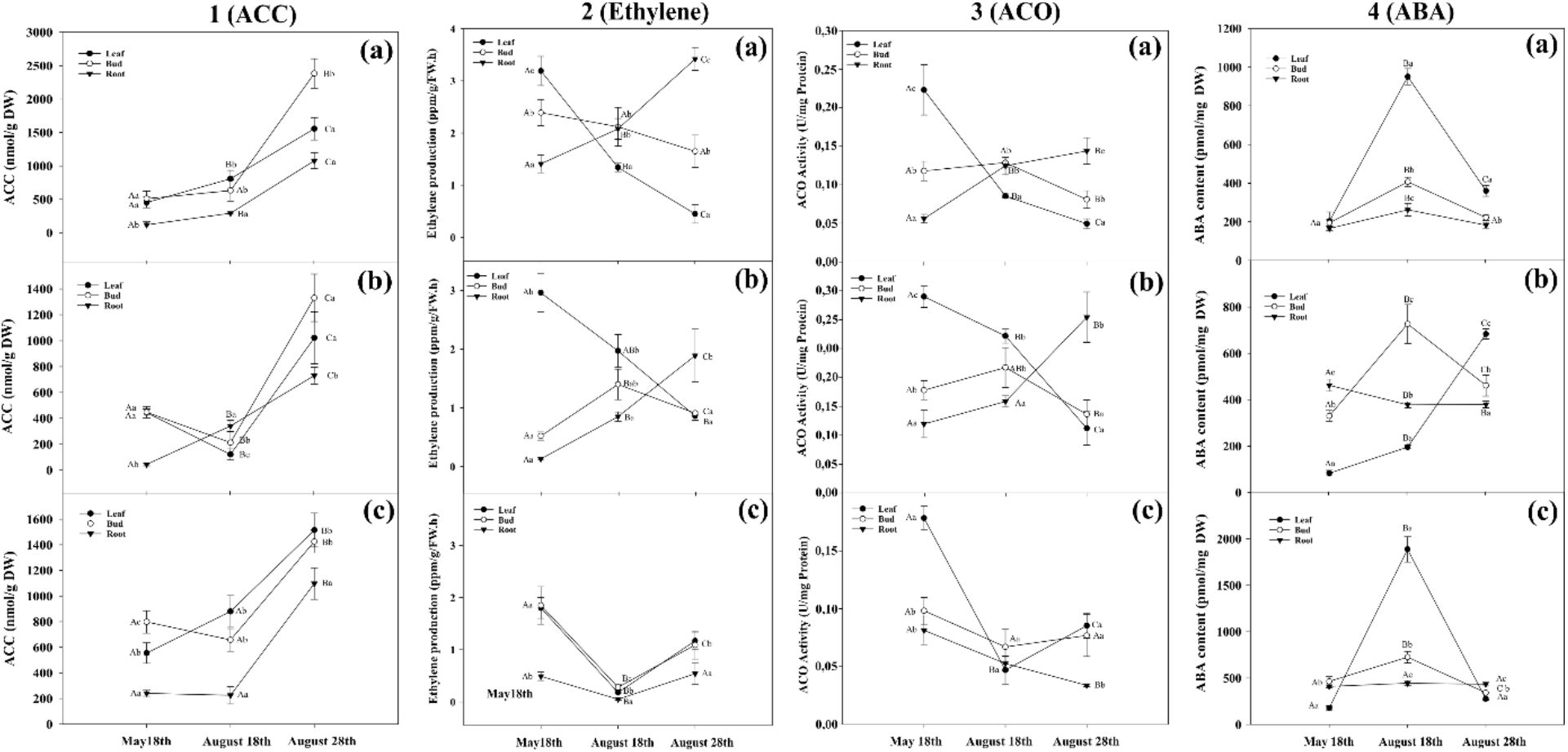
Plant hormones evolution for coffee genotypes. ACC content (**1**), Ethylene production (**2**), ACO activity (**3**) and ABA content (**4**) in leaves, flower buds, and roots from (**a**) Oeiras, (**b**) Acauã, and (**c**) *Semperflorens* plants at the rainy period (May 18th), at the dry period (August 18th), and after rain in the re-watering period (August 28th). Data are means ± 95 % Standard Deviation (SD) of the mean (n = 6). Different upper-case and lower-case letters indicate the statistical difference of each tissue among the different periods and different tissues within the same period, respectively.

### 2.4 Physiological and biochemical analysis

#### 2.4.1. ACC quantification

Leaf, flower bud, and root ACC concentrations were determined by the Bulens method (Bulens *et al*., 2011), with minor modifications. Coffee tissues were grinded in liquid nitrogen using a mortar and pestle. 200 mg were transferred to 2 mL tubes for extraction with 1 mL of sulfosalicylic acid 5 % (p/v). After homogenization, tubes were maintained at 4 ºC for 30 min, being gently mixed with 5 min intervals. After centrifugation at 3,090 g and 4 ºC for 30 min, 300 μL of the supernatant was transferred, in duplicates for each sample, to 10 mL vacutainer glass followed by 100 μL of 10 mM HgCl_2_. The tubes were subsequently sealed with a serum cap, and 300 μL of cold NaOH-NaOCl solution (NaOH 0.434 M and NaOCl 0.173 M) was added using a syringe. The mixture was then homogenized for 5 s and incubated for 4 min on ice. After mixing the samples for 5 s, 5 mL of the headspace was withdrawn and ethylene levels were measured using the F-900 Portable Ethylene Analyzer (Felix Instrument USA) operating under the GC emulation mode. The ethylene ppm values were transformed to nmol using the gases equation, and the ACC content was expressed as nmol/g Dry Weight (DW).

#### 2.4.2. ACO enzymatic activity quantification

Leaf, flower bud, and root ACO activity were determined by the Bulens method (Bulens *et al*., (2011), with some modifications. After tissue grinding in liquid nitrogen, 250 mg were transferred to 2 mL tubes with 50 mg of polyvinylpolypyrrolidone. ACO extraction was performed with 1 mL of extraction buffer (MOPS 400 mM pH 7.2; 30 mM of ascorbic acid, and glycerol 10% (v/v)). Subsequently, samples were homogenized and centrifuged at 22,000 g for 30 minutes. Then, 800 μL of supernatant was collected and used for the subsequent analysis. 400 μL of the enzymatic extract was added to 3.6 mL of a reaction buffer containing: 2.8 mL of MOPS buffer (MOPS 64.3 mM pH 7.2; glycerol 12.86 % (v/v); sodium bicarbonate 25.8 mM; and iron sulfate 26 μM), 0.4 ml of ascorbic acid 45 mM, 0.1 mL of ACC 36 mM, and 0.3 mL of dithiothreitol (DDT) 12 mM. After sample homogenization, the reaction for ethylene release was carried out for 20 min at 30 °C. After homogenization by 5 s, 5 mL of the headspace air was withdrawn and used for the ethylene measurements using the F-900 Portable Ethylene Analyzer (Felix Instruments, USA) operating under the GC emulation mode. The ethylene ppm values were transformed to nmol using the gases equation, and the protein content was determined by the Bradford method (Bradford, 1976) in duplicate, using bovine serum albumin (BSA) as standard. One unit of ACO activity was defined as 1 nmol of ACC converted to 1 nmol of ethylene per min at 30 °C (Dong *et al*., 1992).

#### 2.4.3. Ethylene measurement

For all sampling periods, leaves, flower buds, and root tissues were immediately incubated in 10 mL vacutainer glass tubes, containing a moist tissue placed on the bottom of each vial, sealed with serum caps, and incubated for 24 hours. For each biological sample, ethylene was quantified from the headspace gas using the F-900 Portable Ethylene Analyzer (Felix Instruments, USA) operating under the GC emulation mode in triplicate. Plant material was incubated in two separate vials, and the headspace gas was withdrawn from the vials with a 10 mL plastic syringe. Samples, made of 2.5 mL of gas from each vial, were extracted using the same syringe and subsequently injected into the Ethylene Analyzer. After ethylene measurement, plant material was weighed and ethylene production rate was expressed as ppm/g FW. h.^-1^ (Fresh weight per hour)

#### 2.4.4. ABA extraction and concentration measurement

ABA content analysis in leaves, flower buds, and roots was carried out according to Liu *et al*. (2014). Samples were grinded in liquid nitrogen, then 500 μL of an extraction solution (methanol 90 % (v/v) and sodium diethyldithiocarbamate trihydrate 200 mg/L) was added to 200 mg of plant material in a siliconized borosilicate tube. Samples were incubated overnight under dark conditions at 4 °C and centrifuged at 8,000 g for 10 min at 4 °C. The supernatant was transferred to a pre-cold 1.5 mL Eppendorf tube and evaporated in a vacuum centrifuge at room temperature. The residue was dissolved in a methanolic Tris buffer solution containing methanol 10 % (v/v), Tris-HCl 50 mM pH 8.0, MgCl2 1 mM, and NaCl 150 mM. The ABA concentration in all tissues was measured with a Phytodetek ABA enzyme immunoassay test kit (Agdia; Elkhart, IN, USA, Catalog number: PDK 09347/0096), in duplicate according to the manufacturer’s instructions.

### 2.5 Molecular analysis

#### 2.5.1. *In silico* Analysis

Genes encoding for Lysine Histidine Transporter (LHT1) in *Arabidopsis thaliana* were retrieved from the Arabidopsis Information Resource (https://www.arabidopsis.org/index.jsp) database. The protein sequences from these genes were used as input to perform similarity searches against the genomes of plant species from different orders such as *Rubiaceae, Solanales, Rosales, Gentianales, Malpighiales, Vitales, Poales*, and *Amborellale* by the Protein Basic Local Alignment Tool (BLASTp), at National Center for Biotechnology Information (NCBI, http://www.ncbi.nlm.nih.gov/). The sequences with significant similarity (e-value <10−5) were selected and the predicted proteins in which the inputted sequence identity was below 70% were removed. Protein sequences were aligned using the Clustal W program (Thompson *et al*., 1994) with the standard patterns. The phylogenetic tree was drawn using the MEGA software version 6.0 (Tamura et al., 2013), with a neighbor-joining comparison model (Saitou et al., 1987) and bootstrap values from 5,000 replicate to assess the robustness of the tree. The *LHT1*primer (Forward 5’TTCGTCGGTTGCTCATCTCA and reverse: 5’TTGCCTTCTTCAGCCGTT) design was performed using the sequence obtained by the *in silico* analysis and the Primer Express v2.0 program (Applied Biosystems) (Supporting information Table S2).

#### 2.5.2. RNA extraction, cDNA synthesis, and RT-qPCR assay

Total RNA from leaves, floral bud, and roots were extracted according to de Oliveira *et al*. (2015), with minor modifications. RNA samples (7,5 μg) were treated with DNase I using the Turbo DNA-free Kit (Ambion) to eliminate DNA contamination. RNA integrity was analyzed in 1 % agarose gel, and RNA content, as well as quality, were accessed by spectroscopy (OD260/280 and OD260/230 *>* 1.8) (NanoVue GE Healthcare, Munich, Germany). One μg of the total RNA was reverse transcribed into cDNA using the High-Capacity cDNA Reverse Transcription Kit (Thermo Fisher Scientific, Waltham, USA), according to the manufacturer’s protocol.

Real-time quantitative PCR (RT-qPCR) was performed using 15 ng of cDNA, with Rotor-Gene SYBR® Green PCR Kit (Qiagen), using a Rotor Gene-Q(R) thermocycler (Venlo, Netherlands). Reactions were carried out in 15 μL total reaction volume: 7.5 μL of SYBR-green (QuantiFast SYBR Green PCR Kit - Qiagen), 0.3 μL of forward and reverse gene-specific primers, 1.5 μL of cDNA at 10 ng/μL, and 5.7 of RNase-DNase-free water. Three biological repetitions were used, and reactions were run in duplicate as technical replicates. Amplification was performed with the following reaction conditions: enzyme activation with 5 min at 95 °C, then 40 cycles of 95 °C for 5 s, followed by 10 s at 60 °C and completed by a melting curve analysis to assess the specificity of the reaction by raising the temperature from 60 to 95 °C, with 1 °C increase in temperature every 5 s. Relative fold differences were calculated based on the ΔΔCT method (Pfaffl, 2001), using *MDH* and *RPL39* as reference genes (De Carvalho *et al*., 2013; Fernandes-Brum *et al*., 2017) (Supporting information Table S3).

### 2.6 Statistical analysis

For ACC, ethylene, ACO activity, and ABA, data analysis was performed using the InfoStat software (Di Rienzo *et al*., 2020). The statistical difference was determined by one-way ANOVA, followed by the Tukey test. Results were expressed as the mean ±Standard Deviation (SD). The values marked with different letters are significantly different at P<0.05. For the gene expression, statistical analyses were performed by the R software (R Core Team, 2019). The expression rate and the confidence intervals were calculated according to the method proposed by Steibel *et al*. (2009), which considers the linear mixed model given by the following equation: yijklm = μ + TGijk + Il + eijklm where, yijklm is the Cq (Quantification cycle) obtained from the thermocycler software for the kth gene (reference or target) from the mth well, corresponding to the lth plant subject to the *i*th treatment (Wet, Dry, and Rainy) at the *jth* tissues (Leaf, Bud, and Root); TGijk is the effect of the combination of the *ith* treatment (May 18th, August 18th, and August 28th) at the *jth* tissues (Leaf, Bud, and Root). Graphics were performed with SigmaPlot v. 14 (Systat Software Inc.).

## 3. RESULTS

### 3.1 Leaf water potential

In field experiment I, leaf water potential was measured from -0.35 to-0.45 MPa in the rainy period (May 18^th^), decreasing dramatically at the end of the dry period (August 18^th^), reaching values from -2.5 to -2.7 MPa. After 22 mm of rain in the re-watering period (August 28^th^), leaf water potential values increased to -1.5 MPa (Fig. 1a). In field experiment II, leaf water potential was measured only in the re-watering period, showing that values were higher in covered than in uncovered plants (Fig. 1b).

### 3.2 Field Experiment I

#### 3.2.1 ACC content

In general, higher ACC levels were observed in the cultivar Oeiras when compared to the other genotypes (1.5 times more than Acauã and *Semperflorens*). Within each cultivar, there were different patterns of ACC production across the periods. For Oeiras, ACC showed an increase along the sampling periods for leaves, flower buds, and roots (Fig. 2 (1a)), whereas for Acauã, there was a decrease in the levels in the dry period for leaves and flower buds and an increase in the root, with increased levels being observed in all tissues after the rain event (Fig. 2 (1b)). As for the cultivar *Semperflorens*, a pattern of ACC increase in the leaves was observed during the dry period, slightly decreasing in flower buds, and remaining stable in roots. Similar to the previous genotypes, ACC levels increased after the rain event for leaves, flower buds, and roots (Fig.2 (1c)).

Although all genotypes presented different patterns of ACC production in their different tissues between the rainy and dry periods, the first rain after the dry period provided increases of ACC observed in all tissues from the three coffee genotypes (Fig.2 (1)). Oeiras ACC levels increased 93%, 277%, and 270% for leaves, flower buds, and roots, respectively, once plants were partially rehydrated. For Acauã plants, ACC also increased greatly in root and leaf (119 % and 713 %, respectively), as well in *Semperflorens*, rain promoted ACC increases of 72 % (leaves) and 388 % (roots) related to the dry period. When comparing the production of ACC among the three genotypes, it could be observed that there is a difference between Acauã and Oeiras where, although they showed similar ACC levels until the end of the dry period, ACC levels after the rain ranged from 800 to 1200 nmol ACC/gr DW (Acauã) and 1500 to 2300 nmol ACC/gr DW (Oeiras) (Fig. 2a and 2b). In *Semperflorens* plants, the pattern of ACC production was similar to the one observed for Acauã plants (Fig. 2 (1c)).

#### 3.2.2 Ethylene production

Ethylene production showed different patterns among tissues and genotypes. Oeiras ethylene production decreased along the sampling periods for leaves and flower buds, with the opposite being observed for roots (Fig. 2 (2a). Acauã ethylene production decreased in leaves and increased for roots along the sampling periods, whereas for flower buds an increase was observed during the dry period, decreasing after the rain in the re-watering period (Fig. 2 (2b)). In *Semperflorens*, ethylene production showed a similar pattern for the three tissues, decreasing during the dry period and increasing once plants were partially rehydrated after the rain event (Fig. 2 (2c)). In general, ethylene production was similar for the three genotypes, showing a decreasing pattern in leaves along the sampling periods and an increasing pattern in roots of Oeiras and Acauã plants (Fig. 2 (2)).

#### 3.2.3 ACO activity

In Acauã and Oeiras the ACO activity had a similar pattern (Fig. 2 (3a and 3b)), decreasing from rainy period (May 18^th^) to re-watering period (August 28^th^) for leaves, and increasing in roots. Meanwhile, flower buds decreased ACO in the dry period (August 18^th^). In *Semperflorens* plants, ACO activity decreased from the rainy period (May 18^th^) to the dry period (August 18^th^) for all tissues and increased in the re-watering period (August 28^th^) for leaves and buds but continued decreasing for roots (Fig. 2 (3c)).

#### 3.2.4 ABA content

ABA content was different among the analyzed sampling periods. In general, it was low in the rainy period (May 18^th^), increasing in the dry period (August 18^th^) and re-watering period (August 28^th^) observing different behaviors dependent on each cultivar. The ABA content was higher in *Semperflorens* than Oeiras and Acauã, especially for leaves. For Oeiras, the ABA level was similar for leaves, buds, and roots in the rainy period (May 18^th^), and showed higher levels for all tissues in the dry period but being even higher in leaves. Furthermore, in the re-watering period (August 28^th^) the ABA level decreased for all tissues, although was higher in leaves compared to other tissues, having levels similar to the dry period (Fig. 2 (4a)). In Acauã leaves ABA content increased from the end of the rainy period to the re-watering period (May 18^th^ to August 28^th^), whereas, buds increased ABA from rainy to dry period (May 18^th^ to August 18^th^) and decreased in the re-watering period (August 28^th^); ABA in roots was high in the rainy period (May 18^th^), and stable during the dry and re-watering periods (August 18^th^ and August 28^th^) (Fig. 2 (4b)). In *Semperflorens* ABA content had a similar pattern to Oeiras, having low levels in the rainy period (May 18^th^), increasing in the dry period (August 18^th^), and decreasing in the re-watering period (August 28^th^) (Fig. 2 (4c)).

### 3.3 Lysine-Histidine Transporter 1 (*LHT1*) gene expression

In field experiment, I, the relative gene expression of the *LHT1* transporter was compared for the same tissues (Leaves, buds, and roots) among sampling periods. In general, differences were observed in the relative gene expression for root and bud tissues in the re-watering period (August 28^th^) when compared to rainy and dry periods (May 18^th^ and August 18^th^). For leaves tissues, no differences were observed between sampling periods (May 18^th^, August 18^th,^ and August 28^th^) (Fig. 3a). In field experiment II, the comparison of the *LHT1* relative gene expression was made among leaves, buds, and roots for covered and uncovered plants in the re-watering period (August 28^th^). Results showed that *LHT1* expression was higher in uncovered plants for the root and flower bud tissues, whereas, no differences in the relative gene expression were observed in leaves (Fig. 3b), similarly to what was observed in leaves for field experiment I.

**Fig. 3.**
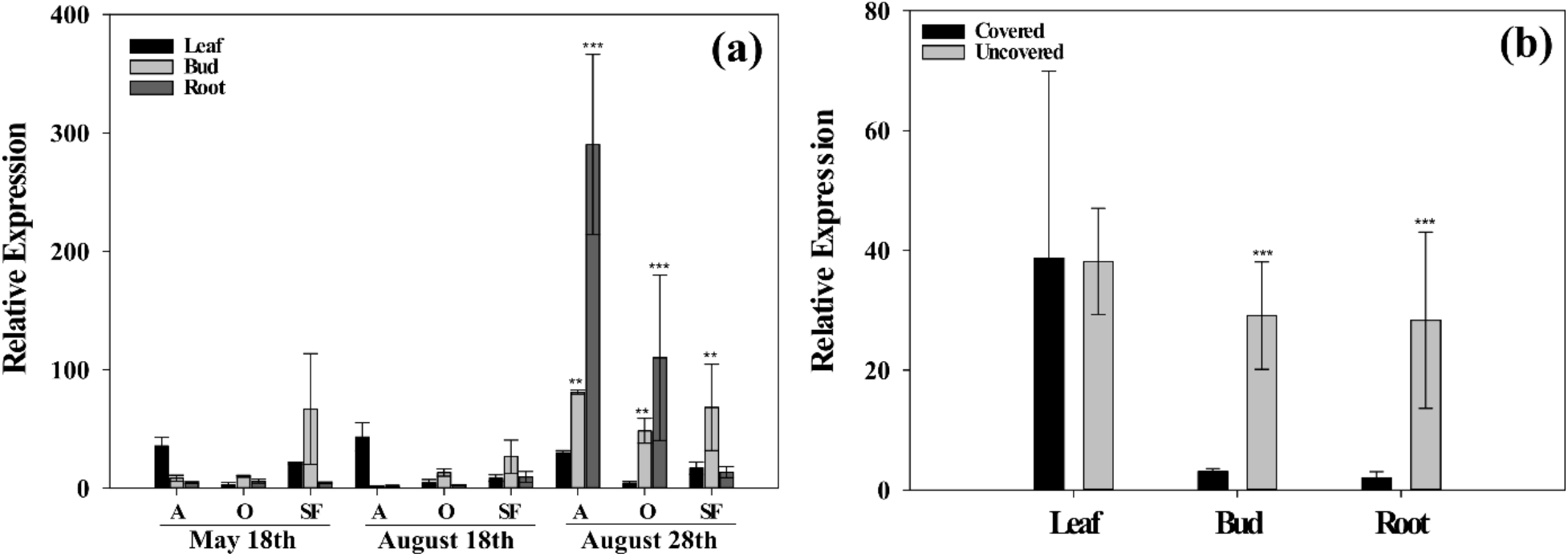
Relative expression of the *LHT1*. (**a**) Field experiment I for leaves, buds, and roots in rainy period (May 18^th^), dry period (August 18^th^), and re-watering period (August 28^th^) for Acauã (A), Oeiras (O), and *Semperflorens* (SF). (**b**) *LHT1* relative expression in leaves, buds, and roots for field experiment II in re-watering period (August 28^th^) for Covered and Uncovered plants of coffee cultivar Paraiso 2. Data are means ± 95 % Standard Deviation (SD) of the mean (n=6). ** = significative difference of tissues between periods and *** = high significant difference of tissues between periods.

### 3.4 Field experiment II

The objective of field experiment II was to evaluate the effect of water on the evolution of ABA, ACC, ACO, and ethylene in the re-watering period (August 28^th^) using two treatments (Covered and uncovered plants). The content of ACC was higher in uncovered plants than in covered ones (Fig. 4a). However, the ethylene production did not vary between treatments, as well as the ACO activity, except for buds, that had higher production and activity in both parameters for uncovered plants (Fig. 4b and Fig. 4c). On the other hand, the ABA content decreased in the uncovered plants (Fig. 4d), possibly as an effect of the rehydration of the plant and the increase in the leaf water potential (Fig. 1b).

**Fig. 4.**
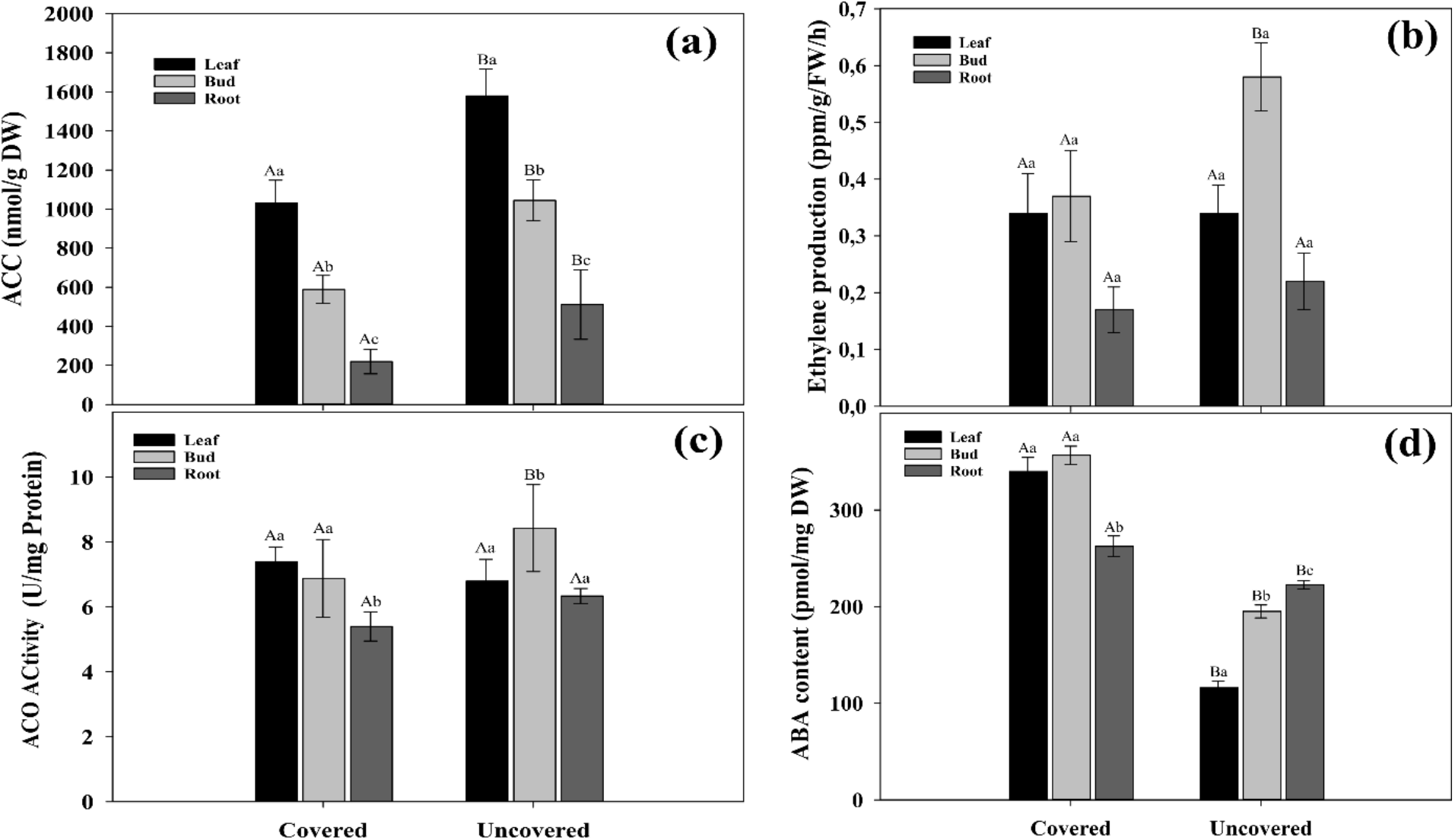
Phytohormones contents. (**a**) ACC, (**b**) Ethylene, (**c**) ACO, and (**d**) ABA quantification in leaves, flower buds, and roots in the field experiment II (covered and uncovered plants of Paraiso 2 coffee cultivar). Data are means ± 95 % Standard Deviation (SD) of the mean (n = 6). Different upper-case and lower-case letters indicate statistical differences of each tissue among the different treatments and the different tissues within the same treatment, respectively.

### 3.5 Field Experiment III

Coffee plants that had exogenous application of ACC showed an increase in the number of flower buds (at the G6 stage). The number of flower buds at the G6 stage was higher in the treatment with ACC (385%), followed by the treatment with 1-MCP + ACC (242%), and 1-MCP (185%) compared with control treatment (Fig. 6a). These results are congruent with the statistical differences found among treatments for flower buds atG6 stage (Fig. 6b). Because all coffee plants had after three days of treatments imposition, a 35 mm rain, the number of flowers bud at G6 stage in control treatment represent the effect of rain in coffee anthesis and the differences between control and the other treatments ((ACC), (1-MCP+ACC) and (1-MCP)) represent the effect those treatments on coffee anthesis.

## 4. DISCUSSION

### 4.1. Water stress and flowering

The transition from the vegetative to reproductive phase is marked by different endogenous processes that respond to environmental stimuli. Diverse stress factors can induce, accelerate, inhibit or delay flowering in many species of plant including water deficit stress (Takeno, 2016). For field experiment, I, Oeiras, Acauã, and *Semperflorens* genotypes were under water deficit stress and leaf water potential reaching -2.5 MPa during the dry period (August 18^th^) and then the average value of - 1.5 MPa after rain in the re-watering period (August 28^th^) (Fig.1a) with anthesis occurring 10 days after 22 mm of rain (Fig. 5a). For field experiment II, covered and uncovered plants showed differences in leaf water potential (Fig. 1b), with only uncovered plants blossoming. (Fig. 5b)

**Fig. 5.**
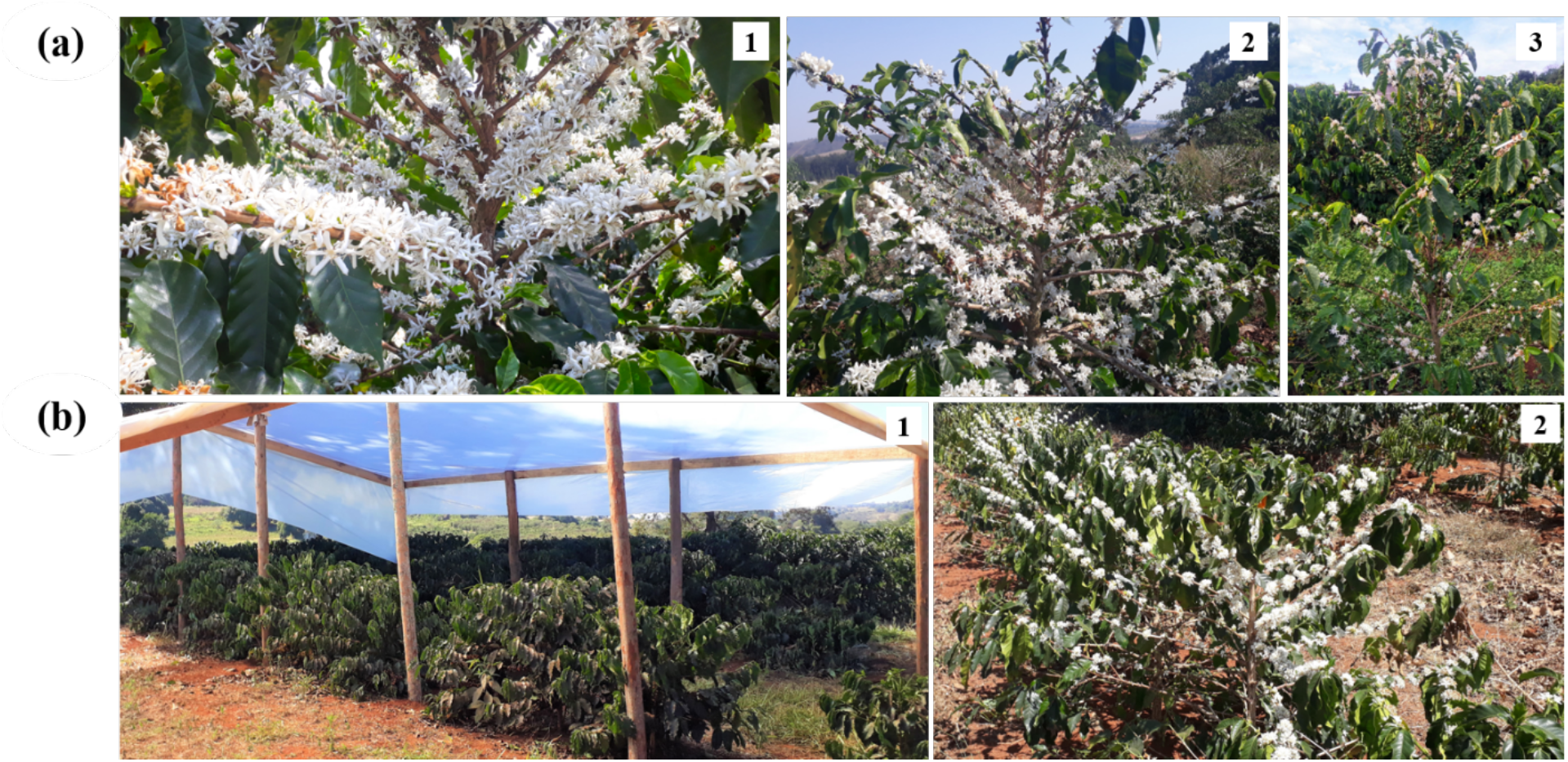
Coffee anthesis in (**a**) Oeiras (1), Acauã (2), and *Semperflorens* (3) in field experiment I. (**b**) Non – Flowering (1) and flowering (2) plants of Paraiso 2 coffee cultivar using treatments of Covered and Uncovered environment at the field experiment II.

In the agroecological conditions of Brazil, coffee anthesis occurs after a short rainy event preceded by a water deficit period during the winter, which is necessary to break the bud “latent state”, associated with endogenous and external factors (Ronchi & Miranda, 2020). Stress inducing flowering can also be seen in lemon (*Citrus limon* L. Burm. f.) under a temperature range from 18 till 30 °C, where flowering was controlled by drought (Chaikiattiyos *et al*., 1994), as well as for “Tahiti” lime (*Citrus latifolia* Tanaka) (Southwick & Davenport, 1986). In Litchi (*Litchi chinensis* Sonn) the autumnal water stress period increased significantly flowering intensity and yield (Stern *et al*., 1993), such as in Star fruit (*Averrhoa carambola* L.) (Wu *et al*., 2017), Longan (*Dimocarpus longan* Lour.) (Zhou *et al*., 2014), and Coffee (*Coffea arabica* L.) (Crisosto *et al*., 1992).

One of the most important factors that contribute to flowering under water deficit stress conditions are phytohormones (Izawa, 2021). The main plant hormone studied in response to water stress is ABA, because acts as an important signaling mediator for plants adaptive response to a variety of environmental stresses regulating many physiological processes, including bud dormancy, seed germination, stomatal development, and transcriptional and post-transcriptional regulation of stress-responsive gene expression (Ali *et al*., 2020).

### 4.2. ABA-Ethylene regulation

ABA and ethylene participate mainly in plant growth regulation, biotic and abiotic stress responses, bud and seed dormancy, leaf and flower senescence, fruit ripening, germination, and flowering (Binder, 2020; Chen *et al*., 2020). In the field experiment I, basically ABA content was higher in leaves and flower buds than in root in the dry period for all genotypes (August 18^th^) (Fig.2(4)), probably because ABA is very active in leaves and buds due to water deficit to keep stomatal conductance in leaves (McAdam & Brodribb, 2018). Conversely, ethylene production decreased (Fig. 2 (2)) coinciding with a reduction of ACO activity, which serves as a catalyzerin the oxidative process for the conversion of ACC into ethylene. This antagonism in response to water stress has been observed in other species directly related to stomata closing to avoid dehydration (Daszkowska-Golec & Szarejko, 2013).

Concerning flowering is known that ethylene could act negatively by inhibiting flowering in *Arabidopsis* (Achard *et al*., 2007) or delaying in rice (Wang *et al*., 2013), demonstrating that ethylene signaling is delayed in both species. However, a positive effect on promoting flowering has been observed in pineapple (*Ananas comusus* L. Merr) (Trusov & Botella, 2006) and lilies (*Triteleia laxa* Benth) (Han *et al*., 1988). Recently, it was proposed that ethylene is involved directly in coffee anthesis by changes in the biosynthesis pathway and regulatory genes expression (Lima *et al*., 2020). In other words, the direct effect of ABA in flowering is not well understood, despite there being some reports about positive and negative influences on flowering (Izawa, 2021). Exogenous application of ABA negatively regulates flowering in *A. thaliana*, represented by *AtABI5* overexpression delaying floral transition by upregulating *FLOWERING LOCUS C (FLC)* expression (Wang *et al*., 2013).

In the same sense, ABA represses flowering by modulating SOC1 at the apex, when CO is needed by FT in the ABA-dependent floral induction (Riboni *et al*., 2020). The most known positive effect of ABA on flowering is the drought escape, which is a plant mechanism to avoid drought damage and reflecting in early flowering, to produce seeds before being affected by severe drought stress conditions (Franks, 2011; Gupta *et al*., 2020). Early flowering is characterized by an ABA increased level in response to water stress (Sherrard & Maherali, 2006; Shavrukov *et al*., 2017).

At the rainy period (May 18^th^), coffee plants showed no water deficit stress (−0.25 MPa) and the ethylene levels were higher than ABA, contrasting with the dry period (August 18^th^), where ethylene level decreased and ABA production increased in coffee plants under water deficit stress (−2.5 MPa) in leaves and flower bud. These results show a clear and opposite behavior in phytohormones content related to water stress conditions in coffee plants (Fig. 2). This behavior suggests a direct ABA regulation on ethylene production, possibly by the downregulation of ACO activity. Negative ABA regulation in the ACO activity was observed in Hibiscus (*Hibiscus rosa-Sinensis* L.) when activation of the ethylene biosynthesis pathway was reduced by exogenous ABA treatment (Trivellini *et al*., 2011). as well as in Lepidio (*Lepidium sativum* L.) when ABA inhibited seed germination decreasing ethylene content by the *Lepidium ACO2* gene expression regulation (Linkies *et al*., 2009). Moreover, in Sugar beet, the radicle emergence is regulated by ABA-ethylene antagonism that affects ACC and ACO gene expression (Hermann *et al*., 2007).

In our research, anthesis occurred after a re-watering period (August 28^th^) by rain, the ethylene content continued to decrease, coinciding with a decrease in the ACO activity. This behavior was observed for leaves and buds, whereas in the roots, it was the opposite. The ABA content decreased in the leaves and flower buds for Oeiras and *Semperflorens* genotypes, possibly because the water content for their rehydration was sufficient enough to reduce the water deficit stress in the plant, whereas in the Acauã cultivar, the ABA content increased in the leaves due to persistent water deficit stress.

In field experiment II, data from ACC, ABA, ethylene, and ACO were taken only after a re-watering period (August 28^th^) to support the previously hypothesized behavior in the coffee plant after rain. The results showed an increased level of ACC and decreased level of ethylene and ACO activity whereas ABA decreased after rain in response to rehydration of coffee plants (Fig. 4). These results corroborate with the results obtained in field experiment I, where an increased level of ACC was observed over an antagonistic relation between ABA and ethylene, participating in coffee anthesis after a water stress period. Moreover, PCA and correlation analysis reinforce the antagonism in the evolution of ABA and ethylene compared in the three sampled periods (Supporting information Fig. S2).

### 4.3. ACC beyond ethylene precursor, acting as a possible flowering signaling

ACC has always been described as an ethylene precursor (Adams & Yang, 1979), and external applications of ACC are used for many experiments with ethylene (Elías *et al*., 2018). However, recent discoveries showed that ACC can act as a signaling molecule in plants, independently of ethylene. One of the most remarkable results of our experiment is the increase of ACC in all tissues and coffee genotypes after rain in the re-watering period (August 28^th^) (Fig. 2 (1)).

This study showed that the increase in ACC found in all tissues does not have a positive correlation with the ethylene production in the coffee plant aerial part (buds and leaves), which coincides with a low activity of the ACO activity in the field experiment I and corroborated with the results in the field experiment II. It is possible that ACC, in addition to being a precursor molecule of ethylene, is possibly involved in the fecundation and anthesis process in coffee. Results of the field experiment III allowed us to confirm this hypothesis because the treatment imposition with exogenous ACC showed a higher number of flower buds at the G6 stage as compared with control treatment (Water) (Fig. 6a, b). As proposed by Lima *et al*., 2020, the 1-MCP application induces ethylene production and participate in the coffee anthesis process. Results of the third field experiment in the treatment where 1-MCP single and combined with ACC was applied, the number of opened and G6 stage flower buds was similar, however in the treatment with only ACC this number was higher. This suggest that, possibly ACC in addition to be acting as ethylene precursor molecule, is directly influencing the coffee flowering anthesis process (Fig. 6a, b). Previous studies in exogenous application of ethylene (Ethrel^®^720) (Data not shown) in coffee flowering effect showed that this phytohormone promotes the senescence process in leaves showing as a yellowing throughout the coffee plant (Supporting information Fig. S3).This high level of ACC observed in all tissues after rain in the re-watering period (August 28^th^) (Fig. 2(1)) could be derived from the amino acid methionine (Precursor of ACC) accumulated during the dry period as a plant response to alleviate the water stress period as observed in previous studies in coffee (Marcheafave *et al*., 2019), wheat (Le Roux *et al*., 2020), and Bitter gourd (Akram *et al*., 2020).

**Fig. 6.**
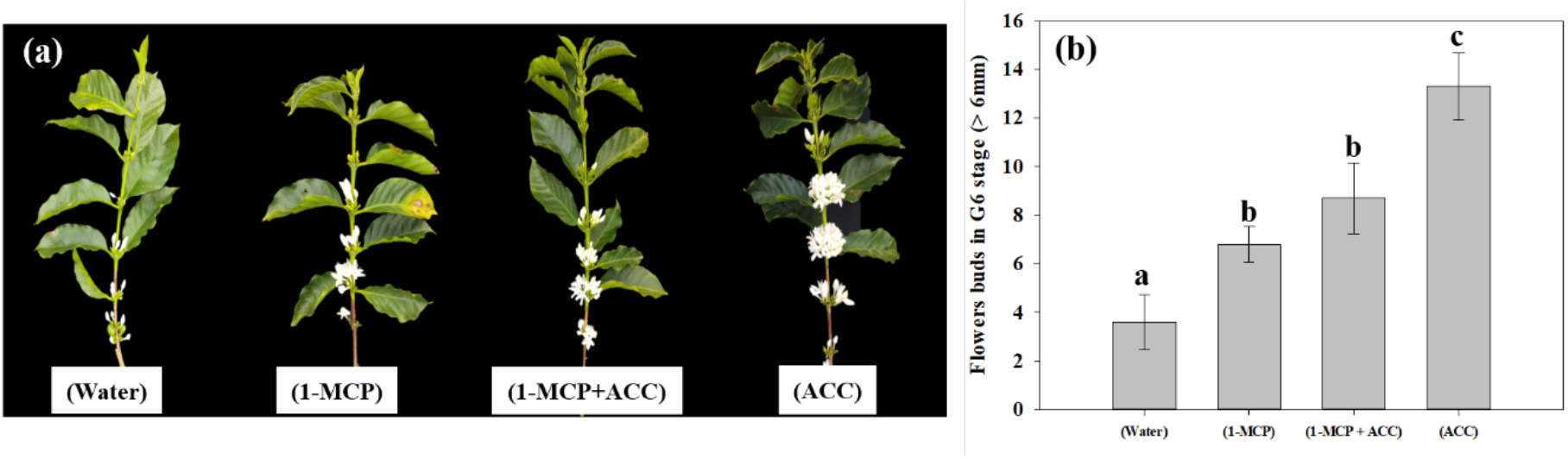
Effect of ACC treatment in coffee anthesis (**a**) Branches with the opened flower as effect different treatments (**b**) Statistical differences among treatments with the application of exogenous ACC and other treatments in flower buds at G6 stage.

In *Arabidopsis*, Mou *et al*. (2020) found that ACC in ovules stimulates transient Ca^2+^ elevation and Ca^2+^ influx, and this signaling in ovular sporophytic tissue is involved in pollen tube attraction. ACC may participate as signaling in pollen tube attraction in the coffee flower fecundation process. This process is very important in the coffee crop because the anthesis only occurs between 10-12 days after the rain, at which time flower fecundation has already occurred at a 90% level because the coffee plant is self-pollinated (Veddeler *et al*., 2006).

ACC has been shown to regulate some physiological mechanisms in plants such as stomatal development by guard cell differentiation (Yin *et al*., 2019), cell wall metabolism (Xu *et al*., 2008; Hofmann, 2008; Tsang *et al*., 2011;), and vegetative development (Tsuchisaka *et al*., 2009; Vanderstraeten *et al*., 2019). In addition, it has been found that ACC has a distinct function than ethylene in the non-vascular plant *Marchantia polymorpha* (Li *et al*., 2020; Katayose *et al*., 2021) and sexual reproduction induction in the Marine red alga *Pyropia yezoensis* (Uji *et al*., 2020). In our study, the *LHT1* expression, an intracellular amino acid transporter that also transports ACC (Shin *et al*., 2015), was evaluated.

In field experiment I, *LHT1* expression was higher in all coffee genotypes after rain in the re-watering period (August 28^th^), especially for roots and buds (Fig. 3a), as well as in the field experiment II, where in the uncovered plants the *LHT1* expression was higher for buds and roots than in covered ones, whereas in leaves it did not show the difference (Fig. 3b). Notably, there was an increase in ACC in the coffee plant tissues (Fig. 2(1)), which coincides with a higher *LHT1* expression after the rain (Fig 4a), showing that LHT1 transporter might be associated with ACC intracellular transportation. The LHT amino acid transporters family are linked with flower development because *LHT2* and *LHT4* were expressed in the tapetum, suggesting their role in delivering amino acids to pollen grains in *Arabidopsis* (Lee *et al*., 2004; Foster *et al*., 2008). Moreover, *LHT5* and *LHT6* expressions were detected along the transmitting tract of the pistil and the pollen tube, pointing to a function in amino acid uptake for successful fertilization (Foster *et al*., 2008). The study represents the first report of *LHT1* transporter gene expression associated with ACC in coffee, suggesting intracellular transport and most likely, participating in the coffee anthesis. In light of these results, we propose that ACC participates in the anthesis process in the coffee plant, as represented in Fig. 7.

**Fig. 7.**
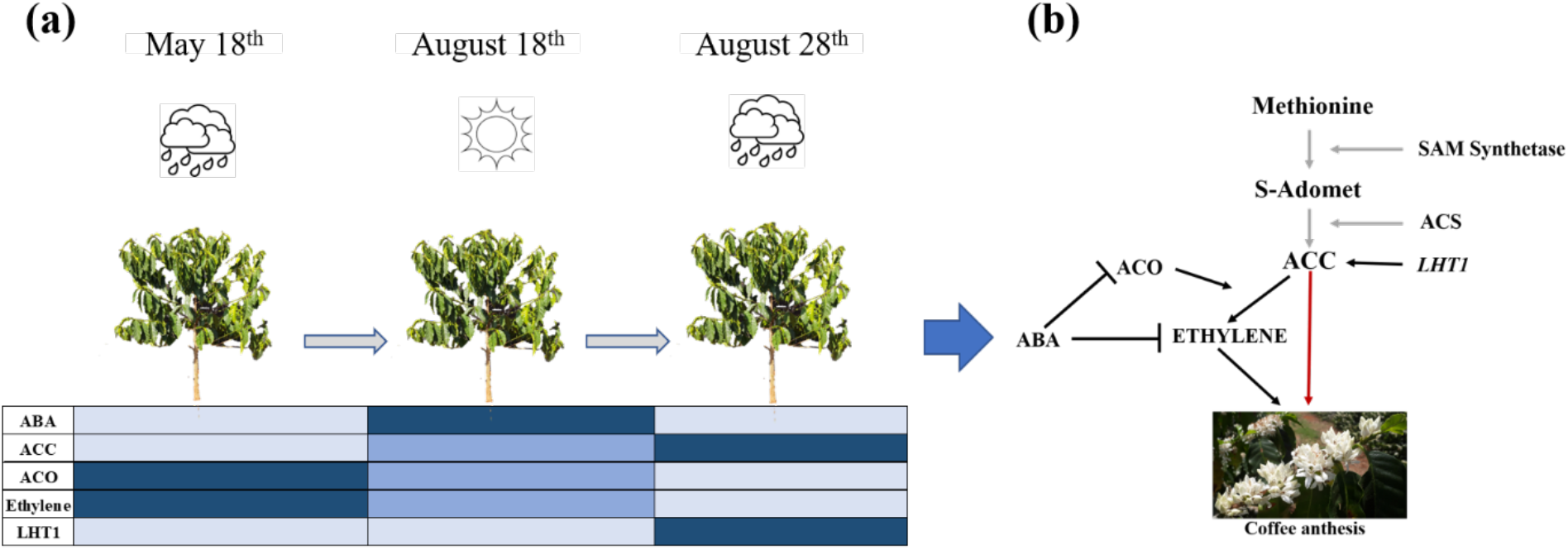
Proposed model of ABA-ACO-Ethylene regulation favoring ACC increased level influencing coffee anthesis in the ethylene biosynthesis pathway. **a**) Evolution of phytohormones (ABA, ACC, ACO, Ethylene) and *LHT1* transporter measured in this study by the sampling times (May 18^th^, August 18^th^, and August 28^th^). The color intensity in the chart means the level of plant hormone in the coffee plant as a whole (Light blue = low, blue = medium, dark blue = high). **b**) Relationship among phytohormones according to our proposal, the black arrow means established plant hormones relation in this study, gray ones mean part of ethylene biosynthesis process described in the literature, the red arrow means a proposal of this study of ACC acting in coffee anthesis.

## 5. ACKNOWLEDGEMENTS

We thank the “Coordenação de Aperfeiçoamento de Pessoal de Nível Superior (CAPES)”, Honduran Foundation for Agricultural Research (FHIA), “Programa de Becas Honduras 2020” for the financial support of the first author and also “Instituto Brasileiro de Ciência e Tecnologia do Café” (INCT/Café), under FAPEMIG grant (CAG APQ 03605/17) for financially support the experiments. The company AgroFresh, for kindly providing us the Harvista used in the experiments of this study and the financial support of the second author.

## 6. AUTHOR CONTRIBUTIONS

MEL, ISS, AAL, and ACJ designed the study, MEL, ISS, JME, and RM performed field experiments, MEL, RM, ISS, AJM and AJM performed laboratory analysis. CC curate and analysed the data, MEL drafted the manuscript before all authors contributed to its improvement and agreed on its final content.

