## Supplementary material for "Crosstalk between Ethylene and ABA during changes in soil water content reveals a role of ACC in coffee anthesis regulation": Suplemental data

The following Supporting Information is available for this article:

**Fig. S1** Design of rainfall exclusion treatment in field experiment II

**Fig. S2 Figure** S2 Spearman Correlation (A) and PCA analysis (B) for ACC, ACO, ethylene, and ABA regulation in coffee anthesis.

**Fig. S3** Effect of ethylene application in coffee plants

**Table S1** Experimental design of the different experiments conducted and their specific hypotheses.

**Table S2** Sequence of primers used for RT-qPCR, where: TM = the melting temperature and E = efficiency of the primer annealing.

**Table S3** Quantitative real-time PCR parameters according to the Minimum Information for publication of Quantitative real-time PCR Experiments (MIQE) guidelines derived from Bustin *et al*. (2009).

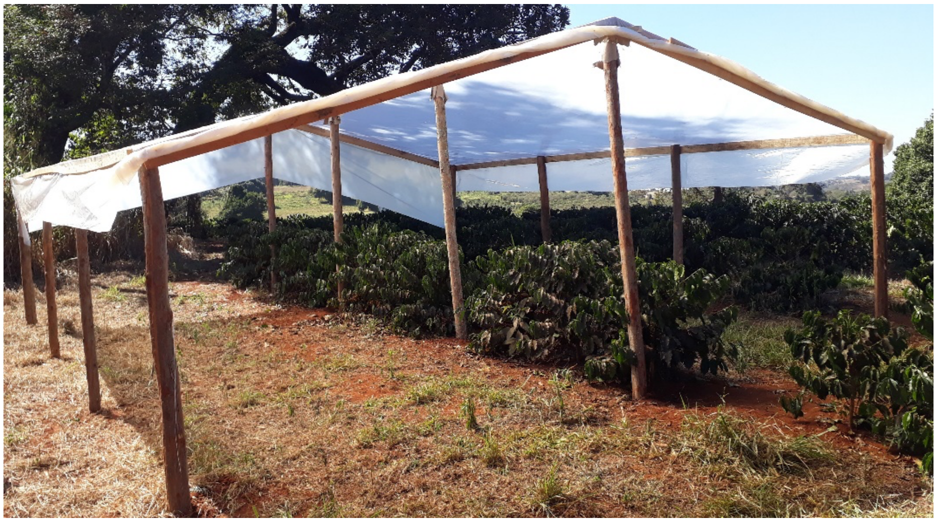

Figure S1: Design of rainfall exclusion treatment in field experiment II. Rainfall exclusion was achieved through the installation of translucent nylon (polypropylene) fixed on twelve 3.2 m high eucalyptus wood logs, buried 0.60 m deep in the soil with three wood logs 3 m distant from each other. Every eucalyptus wood log was arranged on each posterior side and in the middle of the coffee line. A translucent nylon piece of 0,2 micrometers was fixed on top of wood logs using a wooden piece of 5x7 cm with small hobnail, covering a total of 81 $m^2$ of rain exclusion area.

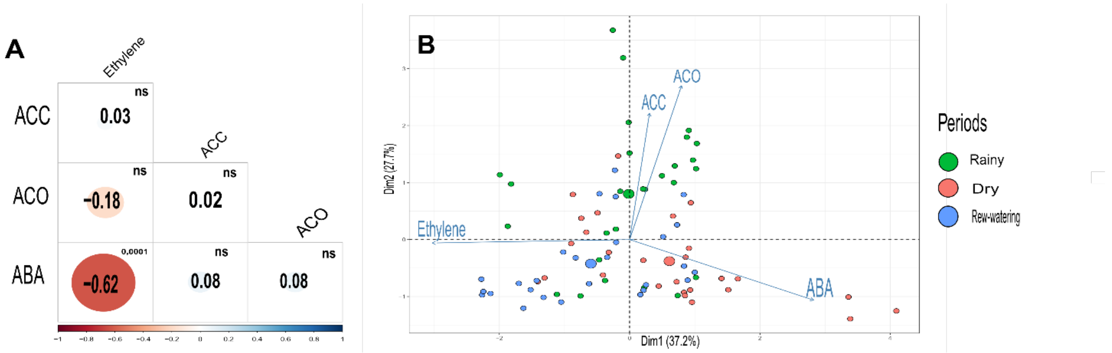


Figure S2: Spearman Correlation (A) and PCA analysis (B) for ACC, ACO, ethylene, and ABA regulation in coffee anthesis. A principal component (PCA) and correlation analysis was carried out to understand the variation and relationship between the variables ABA, ACO, Ethylene, and ACC. The results showed that the biplot PCA represents 64,9% of the total variation. The main component 1 (37,2%) is the ethylene located in the positive part of the axis and by the ABA which is located in the negative part of the X-axis, which indicates that both have a negative correlation. The principal component 2 (27,7%) is represented by the variables ACO and ACC, which are located on the positive part of the Y-axis and are closely related to each other (Fig. 3B). The results of the antagonistic relationship between ABA and ethylene are corroborated by the Spearman correlation analysis where its coefficient is r = -0.62, showing a high negative correlation for the two variables.

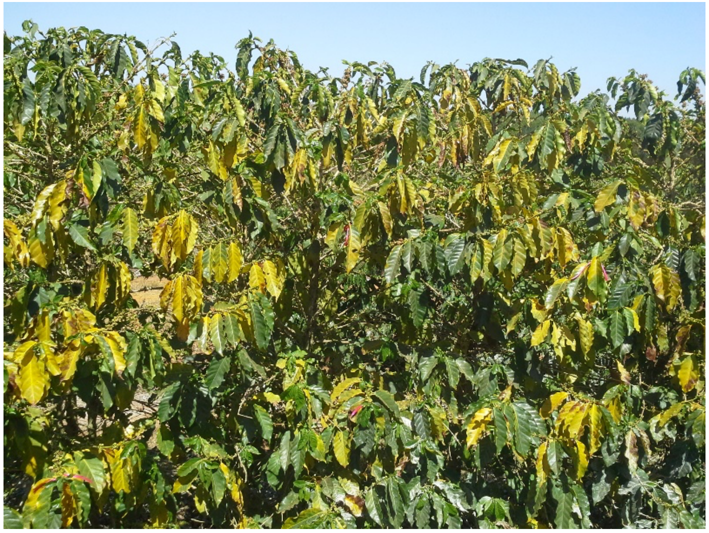

Figure S3 Effect of ethylene application in coffee plants. Commercial product Ethrel 720® was applied in a dose of 1.3 ml diluted in 3L of water, sprayed with the aid of a back spray on 5-year-old coffee trees from cv Catuaí 144. About 24 hours after the spraying significant yellowing and abscission of leaves and fruits was verified.

**Table S1** Experimental design of the different experiments conducted and their specific hypotheses.

| Experiment | Treatments / Condition | Measurements | Hypothesis |
|---|---|---|---|

| | | | |
|---|---|---|---|
| Field I | Three coffee cultivars, Oeiras (Early), Acauã (Late) and *Semperflorens* (continuous) for coffee flowering pattern under field conditions in Rainy, Dry and Re-watering periods | -Predawn Leaf Water Potential<br>- ACC, ABA, Ethylene content, ACO activity and LHT1 gene expression | Hormonal crosstalk is involved in coffee anthesis |
| Field II | Rainfall exclusion, derived in Covered and Uncovered coffee plants in Re-watering period | -Predawn Leaf Water Potential<br>- ACC, ABA, Ethylene content, ACO activity and *LHT1* gene expression | Water is an important triggering transport signalling for coffee anthesis. |
| Field III | In coffee plants of *Semperflorens* were applied ACC, 1-MCP, (ACC+1-MCP) and Water treatments | -Flower bud in G6 stage | Exogenous application of ACC modulates coffee anthesis |

**Table S2** Sequence of primers used for RT-qPCR, where: TM = the melting temperature and E = efficiency of the primer annealing.

| Gene | Sequence of primers (5' – 3') | Concentration (µM) | Volume (µL) | TM (ºC) | E (%) | Reference |
|---|---|---|---|---|---|---|
| Lysine histidine transporter (LHT1) | F: TTCGTCGGTTGCTCATCTCA<br>R: TTGCCTTCTCTTCAGCCGTT | 1,5 | 2,25<br>2,25 | 54<br>52 | 100 | This study |
| Malate dehydrogenase (MDH) | F: CCTGATGTCAACCACGCAACT<br>R: GTGGTTATGAACTCTCCATTCAACC | 2 | 3<br>3 | 59<br>60 | 87 | De Carvalho et al (2013) |
| Large ribosomal subunit 39 (RPL39) | F: GCGAAGAAGCAGAGGCAGAA<br>R: TTGGCATTGTAGCGGATGGT | 2 | 3<br>3 | 59<br>60 | 87 | Fernandes-Brum et al (2017) |

**Table S3** Quantitative real-time PCR parameters according to the Minimum Information for publication of Quantitative real-time PCR Experiments (MIQE) guidelines derived from Bustin *et al.* (2009).

| **Experimental design / Sample** | |
|---|---|
| Experimental group | Four- and five-years old coffee (*Coffea arabica*) plants (Field experiment I, II) respectively. |
| Sample | Leaves, Roots and flower buds for both experiments |
| Sampling procedure | Immediately frozen in liquid nitrogen (roots were first washed and then dried using paper towels) |
| Storage conditions / Time of storage | Freezer -80 °C / 3 weeks |
| **RNA extraction** | |
| Processing procedure | Grinding (mortar and pestle) in liquid nitrogen |
| Method | Organic extraction, according to De Oliveira et al., 2015. |

| | |
|---|---|
| RNA: DNA-free | TURBO DNA-*free*™ Kit (Ambion, Thermo Fisher Scientific)- Catalog number: AM1907 - Design of intron-spanning primers whenever possible |
| Nucleic acid quantification | Spectroscopy (NanoVue GE Healthcare) |
| RNA integrity | Agarose gel (1 %) |
| **Reverse transcription** | |
| Kit | High-Capacity cDNA Reverse Transcription Kit (Applied Biosystems, Thermo Fisher Scientific) - Catalog number: 4368814 |
| Reaction conditions | 25 °C 10' / 37 °C 120' / 85 °C 5' |
| Reverse transcriptase | MultiScrib™ MuLV |
| Amount of RNA / Reaction volume | 1 µg / 20 µL |
| Priming strategy | Random primers |
| Storage conditions of cDNA | Freezer -20 °C |
| **RT-qPCR** | |
| Target information | Table S2 |
| Reaction conditions | As stated in the Materials and Methods section |
| *In silico* | Primers were blasted using the BLAST tool at https://www.ncbi.nlm.nih.gov/ |
| Empirical | Primer concentration of 1µM (final concentration on the reaction)<br>Annealing temperature: 60 °C |
| PCR efficiency | 5-fold dilution series of a mixed sample over at least five dilution points and verified to be higher than 80% (E = 10−1/slope). |
| Linear dynamic range | Samples are situated within the range of the efficiency curves for each primer |
| No template control (NTC) | Cq and dissociation curve verification |
| **Data analysis** | |
| Specialist software | Qiagen Rotor Gene-Q Series software (version 1.7) |
| Normalization | Two reference genes / $\Delta\Delta C_T$ method (Pfaffl, 2001) |
